# An evaluation of longitudinal *Anopheles stephensi* egg viability and resistance to desiccation over time

**DOI:** 10.1101/2025.06.23.661102

**Authors:** L. Leite, J.N. Samake, F.G. Tadesse, S.R. Irish, E. Dotson, S. Zohdy

## Abstract

*Anopheles stephensi*, a malaria vector in South Asia and parts of the Middle East, has been detected as an invasive species in numerous African countries in recent years. It threatens to increase malaria disease burden and reverse gains made in malaria control and elimination over the past decades on the continent. To halt further expansion, it is critical to understand the biological characteristics that may have facilitated *An. stephensi* range expansion to Africa. In its invasive range, *An. stephensi* larvae have been found to colonize artificial containers, many of which are shared with *Aedes aegypti*. The success of *Ae. aegypti* as an invasive vector is often attributed to the use of artificial containers and the ability of *Ae. aegypti* eggs to remain viable in the absence of water for months. While *An. stephensi* is found in artificial containers, it is unclear whether the eggs can remain viable without water for extended periods. *Anopheles stephensi* eggs were reported to remain viable in soil for up to 12 days in a study done almost 100 years ago, but this work has not been revisited since. Thus, in this present study, we used egg batches (>100 eggs per batch) from two laboratory strains of *An. stephensi* (SDA500 and STE2) from South Asia and one *Ae. aegypti* strain (LVP-IB12) to evaluate 1) whether *An. stephensi* eggs can remain viable like *Ae. aegypti* when egg substrates are completely dried following standard insectary methods for drying out *Aedes aegypti* egg sheets, and 2) assess egg viability duration at varying temperatures (15°C, 20°C, 25°C 30°C, 35°C) when eggs are held on a moistened substrate in a high humidity environment. *Anopheles stephensi* egg viability and subsequent larval survival was observed consistently when moistened egg sheets were held at 15°C in a high humidity (>75% RH) environment for up to 14 days in both strains. *Anopheles stephensi* eggs were not viable when dried following standard insectary *Aedes aegypti* methods for drying out egg sheets, except when the protocol was amended to include a 15°C storage temperature. Though egg viability and larval survival was observed in the amended protocol for SDA500 and STE2 (16% and 21% respectively), it was significantly less than that of LVP-IB12 (83%) and was only observed in the egg batches stored for the shortest timepoint (seven days post egg collection, three days post complete drying of egg sheets). These findings suggest that *An. stephensi* may remain viable if eggs are transported under ideal conditions (15°C and >75% RH) through trade or commerce routes. Thus, the persistence of *An. stephensi* eggs in the absence of water should be considered in programs that engage in surveillance and control of *An. stephensi* in Africa.

## Background

*Anopheles stephensi* is a native malaria vector in South Asia and parts of the Middle East and an emerging malaria vector in Africa, Sri Lanka, and Yemen (1). Its rapid range expansion, particularly in countries across Africa over the last decade, threatens to undermine global efforts to control and eliminate malaria (2, 3). Since the first detection of *An. stephensi* in Djibouti in 2012, this mosquito has been established and shown to remain persistent year-round, and modeling efforts have demonstrated its ability to persist even through seasonal shifts in weather (4-6). Similarly, *An. stephensi* was recently found to be established in eastern Ethiopia after its first detection in 2016 (7), and a recent malaria outbreak during a dry season in Dire Dawa, Ethiopia, was linked to *An. stephensi* (8). Thus, as an invasive species in Africa, it is critical to understand the biological characteristics that may have facilitated *An. stephensi* introduction, invasion, and establishment. In urban areas of eastern Ethiopia, *An. stephensi* larvae occupy artificial containers (9-11), many of which are shared with *Ae. aegypti*, the primary vector of arboviral diseases, and a mosquito species that is well known for the ability of its eggs to resist desiccation, a mechanism hypothesized to have aided in its global invasion (12). Also, the dry-arid and seasonal climate across eastern Ethiopia and the reliance of *Anopheles* spp. larvae on water for growth and development raise critical questions about *An. stephensi* vector bionomics as an invasive container breeding mosquito. While *An. stephensi* is a widely used laboratory model for investigating malaria vector competence (13-16), information about its egg desiccation tolerance or larval survival is sparse. A study from 1926 revealed that in its native habitat in India, *An. stephensi* survived in soil without water for up to 12 days (17). More recently, *An. stephensi* was reported as a desiccation-sensitive mosquito species following its use as a control in an *Aedes* egg desiccation mechanisms study (18). Thus, in this study, we evaluate *An. stephensi* egg desiccation tolerance and viability in two well-established laboratory strains of *An. stephensi* from South Asia (SDA500 and STE2), where the African strain was likely to have originated (19), to examine 1) whether *An. stephensi* eggs can remain viable, similarly to *Ae. aegypti* when egg substrates are completely dried following standard insectary methods for drying out *Aedes aegypti* egg sheets, and 2) assess egg viability duration at varying temperatures (15°C, 20°C, 25°C 30°C, 35°C) when eggs are held on a moistened substrate in a high humidity environment. These data will provide preliminary insight into the conditions that may have facilitated the introduction of the invasive malaria species in Africa.

## Materials and Methods

### Mosquito sources, rearing and egg laying

#### Anopheles stephensi

SDA500 (BEI Resources, MRA-1326) mosquitoes originally from Sind, Pakistan (20), and *An. stephensi* STE2 (BEI Resources, MRA-128) mosquitoes from Delhi, India (21) have been maintained in the MR4 at CDC since 2011 and 2000, respectively. *Aedes aegypti* strain LVP-IB12 (BEI Resources, MRA-735) mosquitoes were also used for comparison. LVP-IB12 was selected from the LIVERPOOL *Aedes aegypti* line which originated in West Africa and has been in colony at CDC since 2007. A cohort of about five hundred of each mosquito strain were reared in a walk-in environmental chamber in an Arthropod Containment Level 2 insectary at the CDC. The environmental conditions of the chamber were around 27°C and 78% humidity with 12:12 light and dark cycles that include 30-minute periods of sunrise/sunset. Female mosquitoes (7-14 days post emergence) were blood-fed on a live rabbit. Three days post-blood feeding oviposition cups were placed inside each cage. Because *An. stephensi* mosquitoes lay their eggs directly into the water, the oviposition cup was a 50 ml cup lined with Whatman filter paper and filled with purified water with a depth of 1.0 cm. Because *Ae. aegypti* mosquitoes lay eggs above the water line on the walls of the container, the *Ae. aegypti* oviposition cup was a 150 ml cup lined with seed germination paper (Anchor Paper Company, SD7615L, St. Paul, MN) and with purified water covering the bottom of the cup (depth 0.50cm). Since the walls are lined with seed germination paper, the eggs were laid on the seed germination paper creating egg sheets. Twenty-four hours later the oviposition cups were removed.

### Egg collection and counts

To compare whether *An. stephensi* eggs can remain viable similarly to those of *Ae. aegypti* when egg substrates are completely dried following standard insectary methods for drying out *Ae. aegypti* egg sheets, *An. stephensi* eggs were rinsed from the egg cup onto moistened seed germination paper and excess water removed using a modified vacuum filtration funnel (Fig. S1A). The purpose of this was to create *An. stephensi* egg sheets with the same substrate as the *Aedes aegypti* egg were laid. To assess the viability of eggs from the two *An. stephensi* strains at various temperatures and times, eggs were collected on Whatman filter papers to create egg sheets using the same modified vacuum filtration funnel (Fig. S1B). It is common practice in insectaries to collect *An. stephensi* eggs on filter paper in this manner until the eggs are hatched 24-48 hours later. For both studies, a photograph was taken of each egg sheet and the eggs were counted using egg counter software (22). Only egg sheets with at least 100 eggs per sheet were used.

### Comparison of *Anopheles stephensi* and *Aedes aegypti* egg viability when egg sheets are dried following standard insectary methods for drying *Aedes aegypti* eggs

Post-egg collection, four egg sheets (>100 eggs per egg sheet) from each *An. stephensi* strain and the comparison *Ae. aegypti* strain (LVP-IB12) were placed in plastic containers (7×13 inches) with secured lids inside a Percival environmental chamber (Percival Scientific I36VL incubator, Perry, IA). The chamber was set to 20°C and 50% humidity with a 12-hour light and 12-hour dark cycle to mimic laboratory protocols and conditions where *Aedes aegyp*ti eggs are dried for colony maintenance purposes (Fig. S1; Fig. S2). A data logger (Onset, HOBO U12-012, Bourne, MA) was placed inside the chamber for temperature, humidity, and light monitoring. After 3 days, each container was opened with a 2 cm gap between the lid and container on one side.

The egg sheets were completely dry 24 hours later. The egg sheets were then removed from the plastic containers placed in sealable plastic bags (4×3 inches) and held in the rearing chamber until their scheduled hatch timepoint (7 days, 14 days, 21 days, and 28-days post egg collection). The first scheduled hatch timepoint was seven days after eggs were collected and three days after egg sheets were dried completely. On the scheduled timepoints, egg sheets were placed in larval rearing pans with 250 ml purified water and 25 mg ground fish diet (Drs. Foster and Smith, Staple Diet, Quality Koi and Goldfish Food, Rhinelander, WI) to allow eggs to hatch and larvae to emerge. Larval rearing pans were placed in a Bahnson environmental chamber (Bahnson Environmental Chamber CCS-300, Clemmons, NC) at the above rearing conditions. Larvae were reared following established insectary protocols (23). At six days post hatch, fourth instar larvae were counted, and larval survival rate was calculated as follows: larval survival rate = [(total number of fourth instar larvae x 100)/ total number of eggs]. The egg viability/larval survivability assay was done in 3 replicates per mosquito strain. After completing the temperature study below, this comparison assay was repeated at 15°C with the other conditions the same as above.

## Comparison of *Anopheles stephensi* egg viability at different temperatures on moistened egg sheets

To investigate various environmental conditions that would support *An. stephensi* eggs to remain viable for extended periods, *An. stephensi* (SDA500 and STE2) eggs were stored in high humidity (> 75%) environmental chambers set at the following temperatures: 15°C, 20°C, 25°C, 30°C, and 35°C. Following egg collection, 20 egg sheets (>100 eggs per egg sheet) from each *An. stephensi* strain (SDA500 and STE2) were randomly assigned to four timepoints (0 day, 7 days, 14 days, 21 days) for each temperature tested. Each egg sheet was placed in a 250ml cup inside a closed plastic container along with a moist paper towel and a data logger (Onset, HOBO U12-012, Bourne, MA) for temperature, humidity, and light monitoring and then was placed in a Percival environmental chamber set at one of the experimental temperatures and with a 12-hour light and 12-hour dark cycle. The paper towel was remoistened weekly, and the assay was conducted in 2 replicates for each *An. stephensi* strain (SDA500 and STE2).

On the scheduled timepoints, the egg sheets were placed in larval rearing pans and emerging larvae were reared as described in previous experiment. After determining the temperature with the highest larval survival rate for each *An. stephensi* strain (SDA500 and STE2), both assays were repeated with that specific temperature in 3 replicates for further comparison between the studied *An. stephensi* strains and the *Ae. aegypti* strain.

### Statistical analysis

All statistical analyses were performed using GraphPad Prism version 10.

## Results

### *Anopheles stephensi* and *Aedes aegypti* egg viability comparison on a dried substrate

When egg sheets were hatched seven days post egg collection (three days after egg sheets were dried completely) at 20°C following the standard method for drying out *Aedes aegypti* eggs, 80% of *Ae. aegypti* (LVP-IB12) eggs hatched and survived to late larval instars, whereas no hatching was observed in *An. stephensi* SDA500 and STE2 across three replicates (Fig. 1; Table S1).

**Figure 1.**
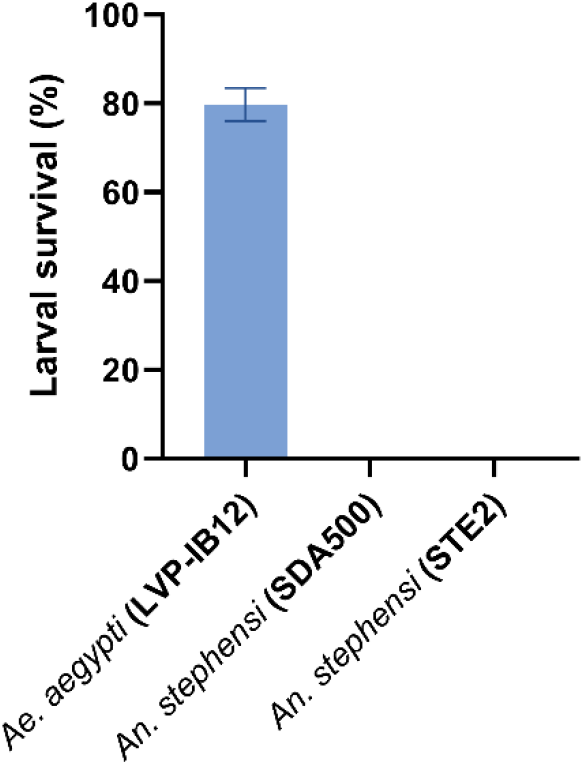
*Aedes aegypti* and *Anopheles stephensi* strains larval survival rates on dried egg sheets stored at 20°C.

### *Anopheles stephensi* egg viability at various temperatures and high humidity

Egg viability and larval survival for both *An. stephensi* strains stored at high humidity (75%) decreased as the number of days post-egg collection increased (Fig. 2). However, larvae from eggs stored at 15°C had the highest larval survival rate at 7 days (SDA500 45%; STE2 58%) and 14 days (SDA500 15%; STE2 4%) post-egg collection for both strains (Fig. 2). No larval survivorship was observed at 21 days post-egg collection across the different temperatures tested (Fig. 2).

**Figure 2.**
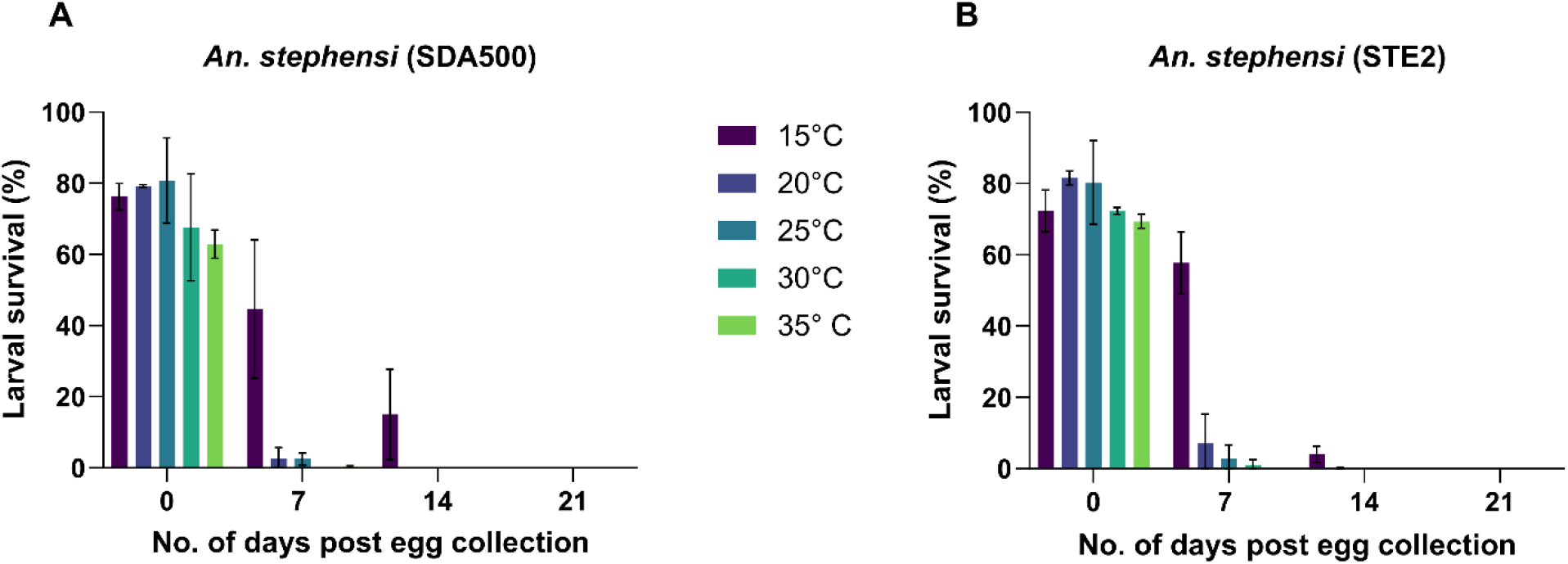
Larval survival rates post egg collection at various temperatures during a period of 21 days in (A) *An. stephensi* SDA500 and (B) *An. stephensi* STE2.

### *Anopheles stephensi* egg viability at 15°C and high humidity

To confirm larval survival when egg sheets are held at 15°C and high humidity, additional replicates were conducted on both *An. stephensi* strains. Eggs viability and larval survival was observed from egg sheets held up to 14 days post-egg collection (Fig. 2). Larval survival rate was the highest for both strains across three replicates at eggs placed in water 0 (SDA500 76%; STE2 77%) followed by 7 (SDA500 31%; STE2 50%), 14 (SDA500 20%; STE2 14%), and 21 days post-egg collection (SDA500 0.6%; STE2 0%) (Fig. 3; Table S3).

**Figure 3.**
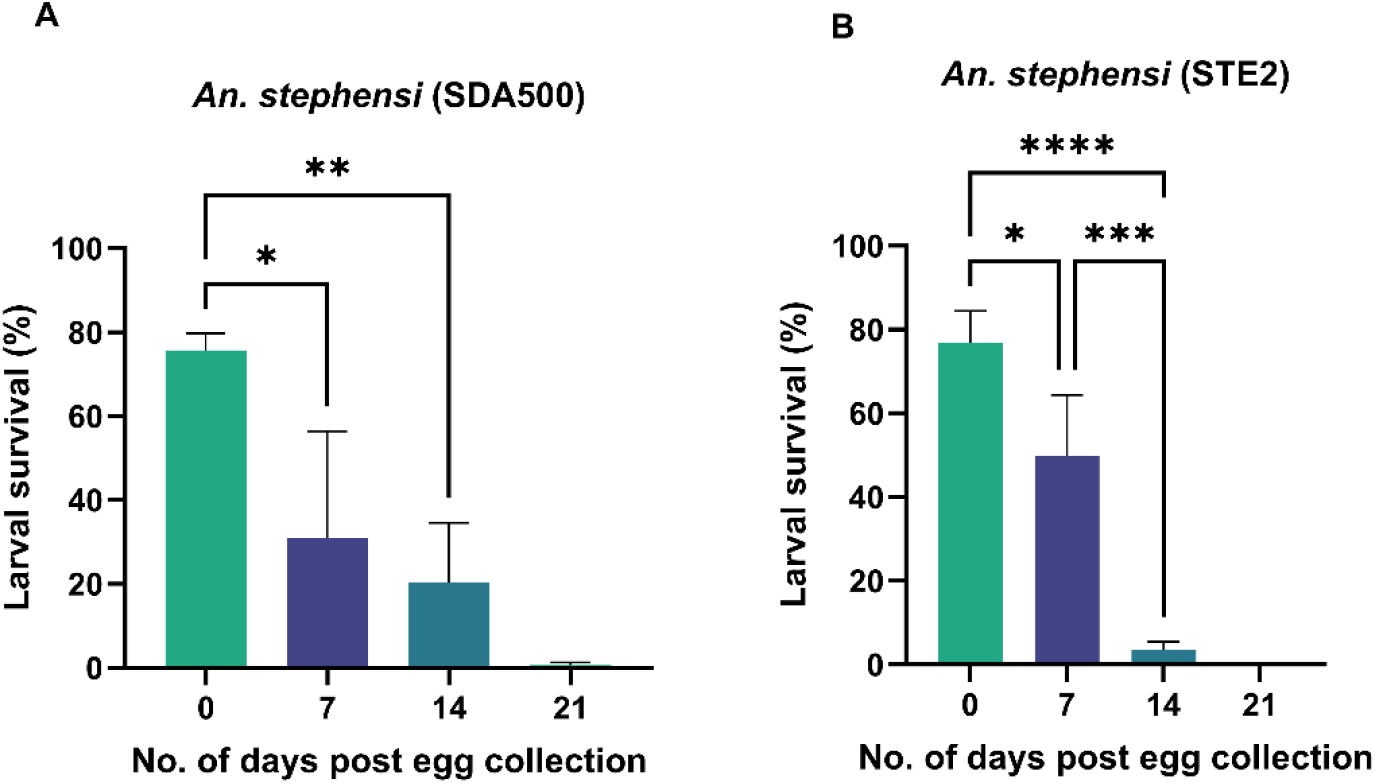
Larval survival rates post-egg collection at 15°C and >75% humidity during a period of 21 days in (A) *An. stephensi* SDA500 and (B) *An. stephensi* STE2.

### *Anopheles stephensi* and *Aedes aegypti* egg viability on dried egg sheets when stored at 15°C

Since *An. stephensi* (SDA500, STE2) eggs stored at 15°C and high humidity on a moistened substrate were observed to be viable for an extended period, the experiment to determine *An. stephensi’s* ability to remain viable when egg sheets are completely dried was repeated with an amended temperature of 15°C (previously 20°C) in two replicates (Table S4). All egg sheets were dried, as previously described. Larval survival rates were 83%, 16% and 21% for *Ae. aegypti* (LVP-IB12), *An. stephensi* SDA500 and STE2, respectively, when eggs were hatched after undergoing the standard method for drying out *Aedes aegypti* eggs at the first scheduled hatch timepoint (i.e., 7 days post egg collection and 3 days after egg sheets were completely dried following the standard method for *Aedes aegypti* egg sheet desiccation) (Fig. 4). When eggs were hatched at the next time point (14 days post egg collection), *Ae. aegypti* (LVP-IB12) had an 87% larval survival rate and *An. stephensi* SDA500 and STE2 both had 0% larval survival rates (Fig. 4).

**Figure 4.**
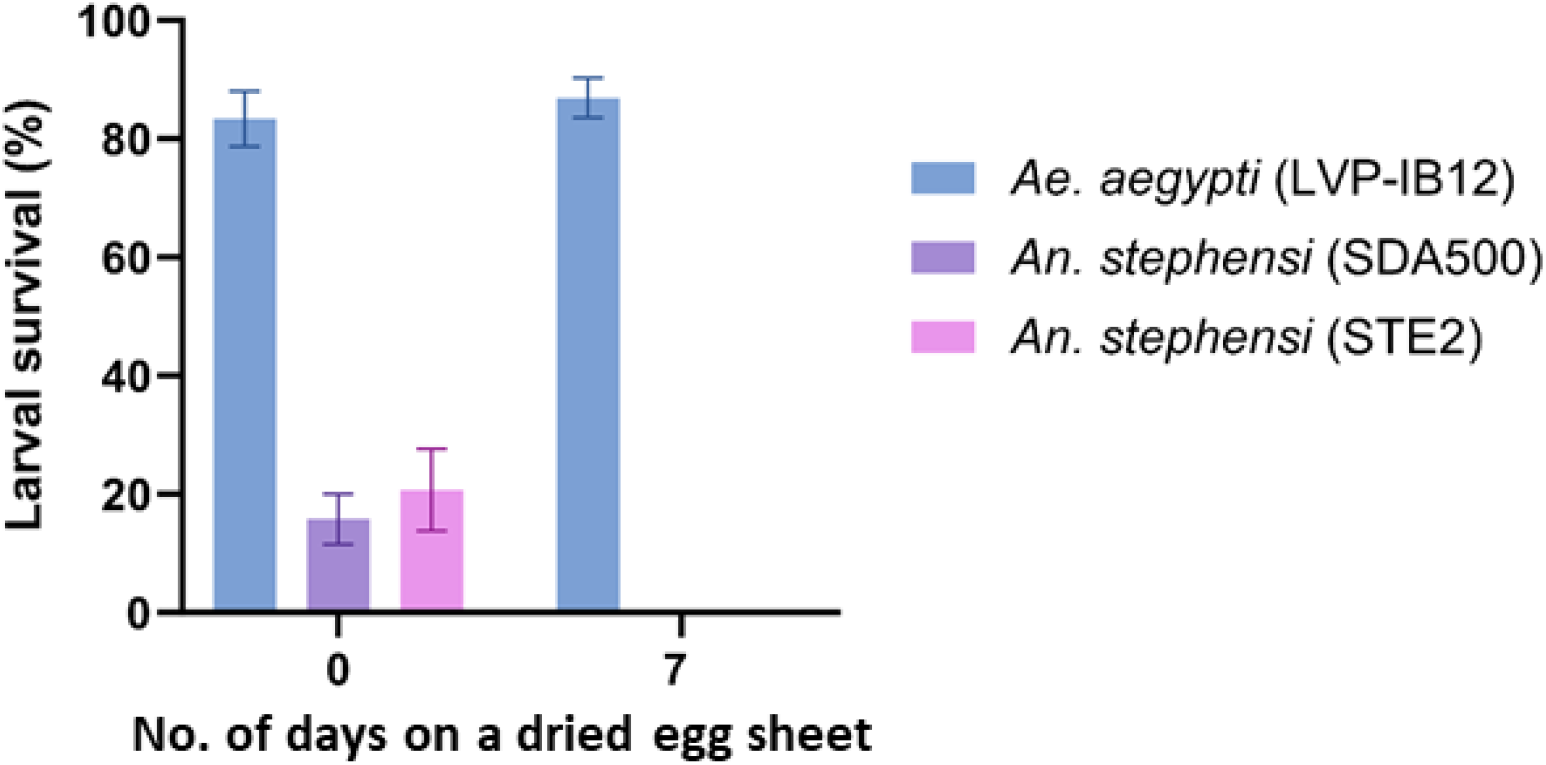
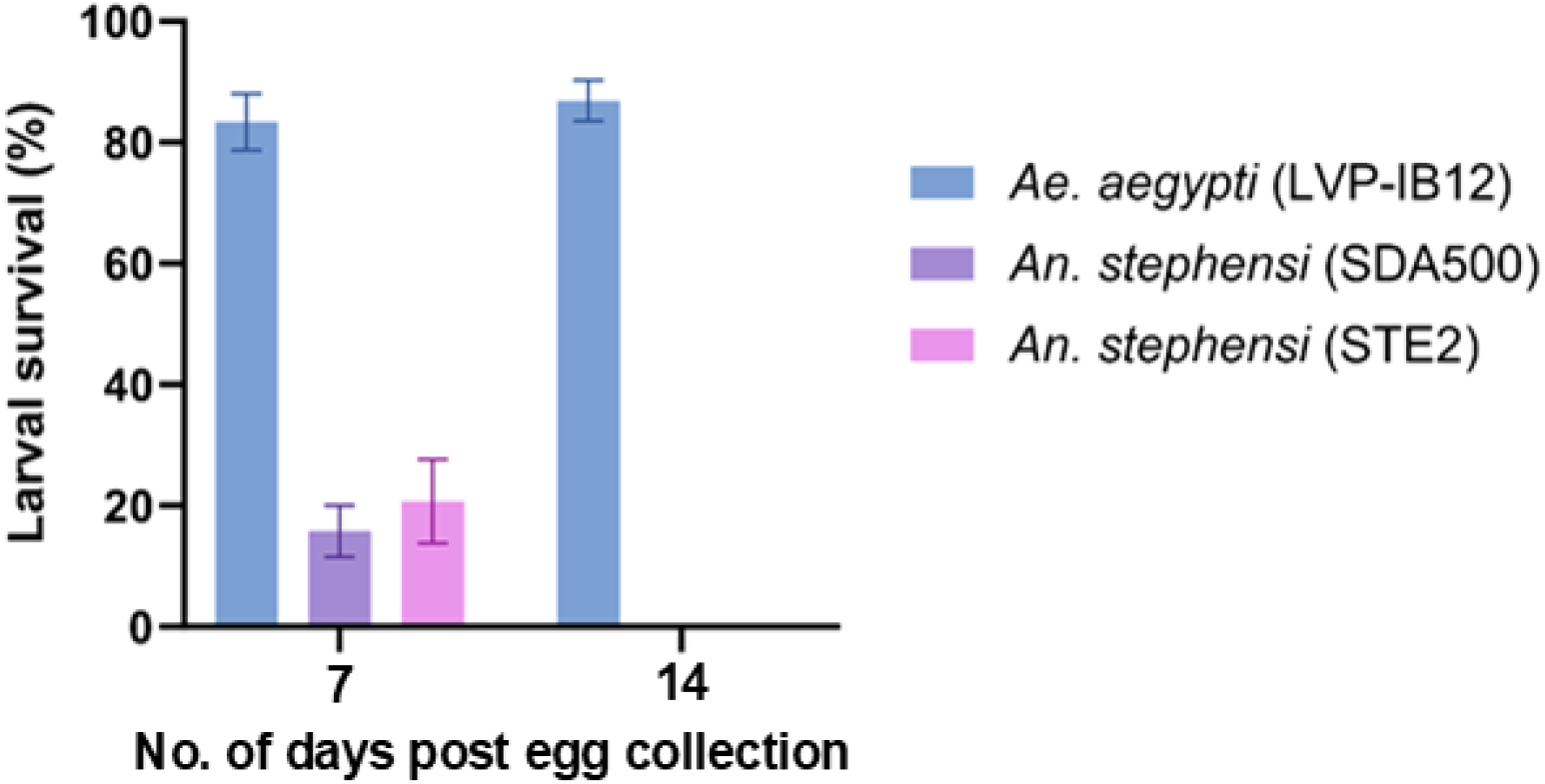
*Aedes aegypti* and *An. stephensi* strains larval survival rates on a dried egg sheet at 15°C.

## Discussion

### Biological Characteristics

Our study revealed that *An. stephensi* (SDA500, STE2) eggs require a moistened substate in a humid environment and an optimum temperature of 15°C for extended egg viability up to 14 days. This finding is consistent with the only previous report of *An. stephensi* egg viability over time, where *An. stephensi* eggs were observed to remain viable in the soil for up to 12 days (17). At the optimum temperature of 15°C, *An. stephensi* (SDA500, STE2) eggs also tolerated being on a completely dried egg sheet, but for a very limited time period. This finding suggests that *An. stephensi* are not completely desiccation-sensitive mosquitoes if given optimal environmental conditions.

### Control Implications

Though *An. stephensi* (SDA500, STE2), unlike *Ae. aegypti* (LVP-IB12), did not remain viable on dry substrates for an extended period, they still showed some level of egg viability under an optimum temperature of 15°C and high humidity. Thus, the persistence of *An. stephensi* eggs in the absence of water should be considered in programs that engage in surveillance and control of *An. stephensi* in Africa. Based on our findings, there may also be opportunities to integrate vector control strategies in urban environments or where *An. stephensi* and *Ae. aegypti* share habitats. Containers of water may be covered to prevent oviposition, emptied when possible, and potentially scrubbed. Scrubbing removes potentially viable eggs from the containers, an action that is more often reserved for the control of *Aedes* (24) but should be considered for *An. stephensi* control as well.

Recent evidence from the Horn of Africa on *An. stephensi* larval habitat characteristics and seasonality has shown that this invasive vector shows the ability to persist through dry seasons and thrive in artificial containers large and small as well as in natural habitats of various sizes (8, 10, 11, 25, 26). Moreover, *An. stephensi* does not follow seasonal population trends as its congeners do (6). However, the underlying mechanisms driving this ecological plasticity remain unknown. Thus, the biological characteristics of *An. stephensi* (SDA500, STE2) egg viability described in this present study may aid in understanding why *An. stephensi* has been a successful container-breeding invasive species.

### Invasion Hypothesis

To control the species and halt further expansion, it is critical to understand the biological characteristics and determinants that may have led to their successful range expansion from South Asia to Africa. Our study showed that *An. stephensi* (SDA500, STE2) eggs can remain viable for several weeks, but only when eggs are on a moistened substrate at 15°C in a high humidity environment. This optimum temperature may offer a possible explanation for their success. Average temperatures in East Africa, the first invaded region in Africa, range between 25°C and 35°C (27). Cargo ships, however, may offer just the right conditions by using temperature-controlled containers for goods transportation between countries. Some containers may provide a cool and humid environment ideal for mosquito eggs, while others may get extremely hot. International trade via import-export connects multiple countries and continents. A commodity often exported and imported worldwide, including in Africa, is rice (28). Rice, similar to wheat and other grains, requires a cool (5°C-20°C) shipping container storage (29) that could also potentially facilitate mosquito egg transport and viability. The role that sea transports and port cities play in facilitating the introduction of invasive mosquito species, particularly *Aedes*, into new geographic areas has been well-documented (30-33). Thus, a similar scenario for the invasion of *An. stephensi* into Africa is plausible. Indeed, most of the first detections of the invasive *An. stephensi* on the African continent have been reported in coastal countries (1).

Moreover, a modeling study predicting countries at risk for *An. stephensi* invasion, based on maritime traffic, accurately identified Sudan and Djibouti with the highest invasion risk (34), as these two countries were among the first to report a *An. stephensi* invasion in Africa (26, 35). The same modeling study also noted that all maritime trade routes could be achieved within 14 days (34). Thus, based on our egg viability findings, a cargo ship carrying *An. stephensi* eggs could reach any port with still viable eggs that could hatch in their new location.

### Limitations

Though our study filled a critical knowledge gap in our understanding of the biological characteristics that may explain the success of *An. stephensi* as an invasive species in Africa, there were some limitations. There was considerable variability in egg counts between egg sheets used in the different assays, which could have impacted the larval survival rate observed. Additionally, eggs were collected onto filter paper using a vacuum platform, as described in the methods section. Leaving the egg sheet on for a shorter/longer duration could have impacted the initial dampness of the filter paper and incorporated variability in the moisture in each assay. A replicate with more initial moisture might avoid drying as quickly and yield a replicate with higher egg viability. Another limitation was the use of filter papers as egg collection and storage substrates. Filter paper is an excellent egg collection substrate but dries out over time, even in high-humidity environments. A substrate that holds moisture longer could be a better alternative.

## Conclusion and future directions

Our present study revealed the biological characteristics that may explain the recent successful *An. stephensi* range expansion to Africa. Eggs from two well-established laboratory strains of *An. stephensi* from South Asia (SDA500 and STE2) remained viable for up to 14 days under an optimum environmental condition of 15°C and >75% RH. Our study also revealed that both *An. stephensi* strains are desiccation tolerant at the same optimum condition (15°C and >75% RH) when held on a moistened substrate, though only for a short period (< 7 days) when the substrate is totally dried. Thus, we recommend that the persistence of *An. stephensi* eggs in the absence of water should be considered in programs that engage in the surveillance and control of *An. stephensi* in Africa. We also recommend that opportunities to integrate vector control strategies in urban environments or where *An. stephensi* and *Ae. aegypti* share habitats should be considered in the invasive range. Overall, this present study provides new insights into *An. stephensi* egg viability and desiccation resistance status over an extended period, which had not been revisited in almost 100 years. However, future studies should evaluate these biological characteristics and environmental conditions in field settings in native and invasive regions to better inform vector control and surveillance strategies to halt the further spread of *An. stephensi* in Africa.

## Disclaimer

The findings and conclusions expressed herein are those of the author(s) and do not necessarily represent the official position of the US Agency for International Development (USAID), the U.S. President’s Malaria Initiative (PMI), or the US Centers for Disease Control and Prevention (CDC).

## Acknowledgements

We would like to thank the MR4 team (Catherine Steele, Rachel Denny, Dustin Miller, Zach Kaplan, Carlye Lapham) for their assistance in providing the mosquitoes used in this study and providing technical advice and assisting with data collection.

The following reagent was obtained through BEI Resources, NIAID, NIH: *Anopheles stephensi*, Strain STE2, MRA-128, contributed by Mark Q. Benedict.

The following reagent was obtained through BEI Resources, NIAID, NIH: *Anopheles stephensi*, Strain SDA-500, Eggs, MRA-1326, contributed by Peter F. Billingsley.

The following reagent was obtained through BEI Resources, NIAID, NIH: *Aedes aegypti*, Strain LVP-IB12, Eggs, MRA-735, contributed by David W. Severson.

