## Supplementary Information for "An evaluation of longitudinal *Anopheles stephensi* egg viability and resistance to desiccation over time"

**A** **B** **C**

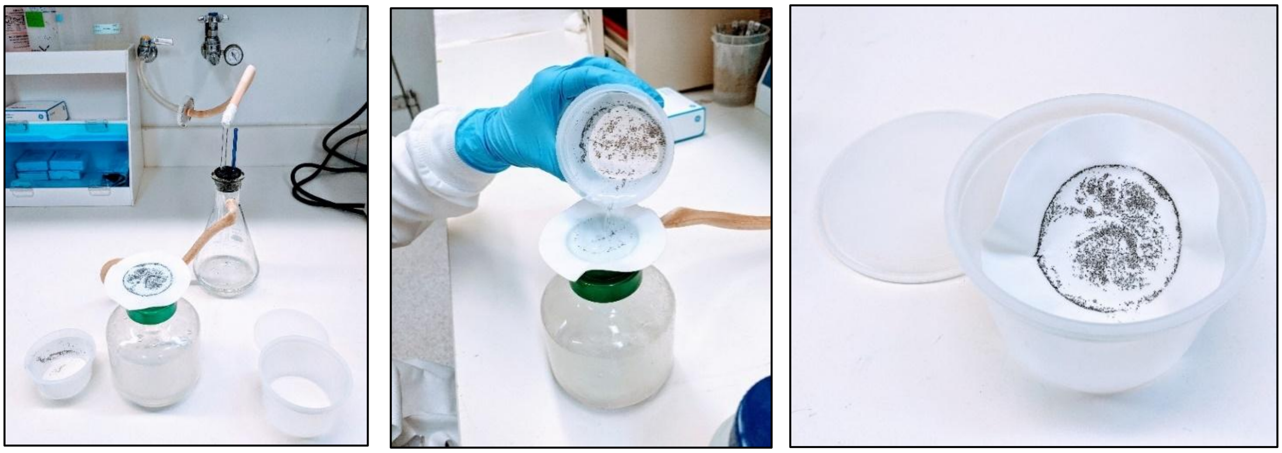

**Figure S1. A-B)** Preparing egg sheets for viability assay. **C)** Egg sheet in a plastic container.

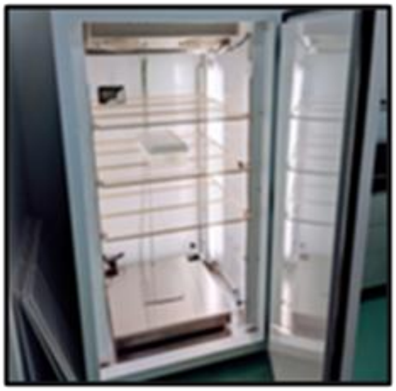

**Figure S2.** Percival environmental chamber.

**Table S1.** *Anopheles stephensi* and *Aedes aegypti* egg viability comparison on a dried egg sheet at 20°C

| Mosquito strain | Replicate | Post egg sheet desiccation | Egg count | Larvae count (L4) | Larval survival (%) |
|---|---|---|---|---|---|
| LVP-IB12 | 1 | 0 | 386 | 309 | 80.1 |
| LVP-IB12 | 2 | 0 | 203 | 154 | 75.9 |
| LVP-IB12 | 3 | 0 | 239 | 199 | 83.3 |
| SDA500 | 1 | 0 | 476 | 0 | 0 |
| SDA500 | 2 | 0 | 221 | 0 | 0 |
| SDA500 | 3 | 0 | 303 | 0 | 0 |
| STE2 | 1 | 0 | 445 | 0 | 0 |
| STE2 | 2 | 0 | 388 | 0 | 0 |
| STE2 | 3 | 0 | 299 | 0 | 0 |

**Table S2.** *Anopheles stephensi* egg viability temperature comparison for (a) SDA500 and (b) STE2.

| **(a) *An. stephensi* SDA500** | | | | | |
|---|---|---|---|---|---|
| **Replicate** | **Temperature (°C)** | **Hatching timepoint (days post collection** | **Egg count** | **Larvae count (L4)** | **Larval survival (%)** |
| 1 | 15 | 0 | 322 | 237 | 73.6 |
| 1 | 20 | 0 | 698 | 551 | 78.9 |
| 1 | 25 | 0 | 374 | 334 | 89.3 |
| 1 | 30 | 0 | 272 | 155 | 57.0 |
| 1 | 35 | 0 | 411 | 247 | 60.1 |
| 2 | 15 | 0 | 133 | 105 | 78.9 |
| 2 | 20 | 0 | 161 | 128 | 79.5 |
| 2 | 25 | 0 | 191 | 138 | 72.3 |
| 2 | 30 | 0 | 184 | 144 | 78.3 |
| 2 | 35 | 0 | 178 | 117 | 65.7 |
| 1 | 15 | 7 | 447 | 261 | 58.4 |
| 1 | 20 | 7 | 429 | 2 | 0.5 |
| 1 | 25 | 7 | 389 | 5 | 1.3 |
| 1 | 30 | 7 | 324 | 0 | 0 |
| 1 | 35 | 7 | 321 | 0 | 0 |
| 2 | 15 | 7 | 175 | 54 | 30.9 |
| 2 | 20 | 7 | 126 | 6 | 4.8 |
| 2 | 25 | 7 | 217 | 8 | 3.7 |
| 2 | 30 | 7 | 139 | 0 | 0 |
| 2 | 35 | 7 | 187 | 1 | 0.5 |
| 1 | 15 | 14 | 429 | 103 | 24.0 |
| 1 | 20 | 14 | 440 | 0 | 0 |
| 1 | 25 | 14 | 372 | 0 | 0 |
| 1 | 30 | 14 | 391 | 0 | 0 |
| 1 | 35 | 14 | 287 | 0 | 0 |
| 2 | 15 | 14 | 168 | 10 | 6.0 |

| | | | | | |
|---|---|---|---|---|---|
| 2 | 20 | 14 | 158 | 0 | 0 |
| 2 | 25 | 14 | 151 | 0 | 0 |
| 2 | 30 | 14 | 188 | 0 | 0 |
| 2 | 35 | 14 | 199 | 0 | 0 |
| 1 | 15 | 21 | 411 | 0 | 0 |
| 1 | 20 | 21 | 368 | 0 | 0 |
| 1 | 25 | 21 | 388 | 0 | 0 |
| 1 | 30 | 21 | 299 | 0 | 0 |
| 1 | 35 | 21 | 262 | 0 | 0 |
| 2 | 15 | 21 | 210 | 0 | 0 |
| 2 | 20 | 21 | 146 | 0 | 0 |
| 2 | 25 | 21 | 133 | 0 | 0 |
| 2 | 30 | 21 | 209 | 0 | 0 |
| 2 | 35 | 21 | 151 | 0 | 0 |
| **(b)** ***An. stephensi*** **STE2** | | | | | |
| **Replicate** | **Temperature (°C)** | **Hatching timepoint (day)** | **Egg count** | **Larvae count (L4)** | **Larval survival (%)** |
| 1 | 15 | 0 | 346 | 236 | 68.2 |
| 1 | 20 | 0 | 368 | 295 | 80.2 |
| 1 | 25 | 0 | 421 | 373 | 88.6 |
| 1 | 30 | 0 | 437 | 313 | 71.6 |
| 1 | 35 | 0 | 448 | 317 | 70.8 |
| 2 | 15 | 0 | 102 | 78 | 76.5 |
| 2 | 20 | 0 | 165 | 137 | 83.0 |
| 2 | 25 | 0 | 186 | 134 | 72.0 |
| 2 | 30 | 0 | 126 | 92 | 73.0 |
| 2 | 35 | 0 | 153 | 104 | 68.0 |
| 1 | 15 | 7 | 327 | 209 | 63.9 |
| 1 | 20 | 7 | 342 | 44 | 12.9 |
| 1 | 25 | 7 | 385 | 21 | 5.5 |
| 1 | 30 | 7 | 373 | 0 | 0 |
| 1 | 35 | 7 | 266 | 0 | 0 |
| 2 | 15 | 7 | 122 | 63 | 51.6 |
| 2 | 20 | 7 | 146 | 2 | 1.4 |
| 2 | 25 | 7 | 232 | 0 | 0 |
| 2 | 30 | 7 | 188 | 4 | 2.1 |
| 2 | 35 | 7 | 146 | 0 | 0 |
| 1 | 15 | 14 | 338 | 19 | 5.6 |
| 1 | 20 | 14 | 388 | 1 | 0.3 |
| 1 | 25 | 14 | 541 | 0 | 0 |
| 1 | 30 | 14 | 266 | 0 | 0 |
| 1 | 35 | 14 | 249 | 0 | 0 |
| 2 | 15 | 14 | 208 | 5 | 2.4 |
| 2 | 20 | 14 | 129 | 0 | 0 |
| 2 | 25 | 14 | 145 | 0 | 0 |
| 2 | 30 | 14 | 201 | 0 | 0 |

| | | | | | |
|---|---|---|---|---|---|
| 2 | 35 | 14 | 178 | 0 | 0 |
| 1 | 15 | 21 | 241 | 0 | 0 |
| 1 | 20 | 21 | 468 | 0 | 0 |
| 1 | 25 | 21 | 323 | 0 | 0 |
| 1 | 30 | 21 | 219 | 0 | 0 |
| 1 | 35 | 21 | 198 | 0 | 0 |
| 2 | 15 | 21 | 150 | 0 | 0 |
| 2 | 20 | 21 | 155 | 0 | 0 |
| 2 | 25 | 21 | 200 | 0 | 0 |
| 2 | 30 | 21 | 207 | 0 | 0 |
| 2 | 35 | 21 | 146 | 0 | 0 |

**Table S3.** *Anopheles stephensi* egg viability at 15°C and high humidity over a period of 21 days for (a) SDA500 and (b) STE2.

| **(a) *An. stephensi* SDA500** | | | | |
|---|---|---|---|---|
| **Replicate** | **Hatching timepoint (day)** | **Egg count** | **Larvae count (L4)** | **Larval survival (%)** |
| 1 | 0 | 322 | 237 | 73.6 |
| 1 | 7 | 447 | 261 | 58.4 |
| 1 | 14 | 429 | 103 | 24.0 |
| 1 | 21 | 411 | 0 | 0.0 |
| 2 | 0 | 624 | 501 | 80.3 |
| 2 | 7 | 570 | 149 | 26.1 |
| 2 | 14 | 944 | 305 | 32.3 |
| 2 | 21 | 846 | 11 | 1.3 |
| 3 | 0 | 764 | 559 | 73.2 |
| 3 | 7 | 602 | 48 | 8.0 |
| 3 | 14 | 743 | 34 | 4.6 |
| 3 | 21 | 483 | 3 | 0.6 |
| **(b) *An. stephensi* STE2** | | | | |
| **Replicate** | **Hatching timepoint (day)** | **Egg count** | **Larvae count (L4)** | **Larval survival (%)** |
| 1 | 0 | 346 | 236 | 68.2 |
| 1 | 7 | 327 | 209 | 63.9 |
| 1 | 14 | 338 | 19 | 5.6 |
| 1 | 21 | 241 | 0 | 0.0 |
| 2 | 0 | 463 | 369 | 79.7 |
| 2 | 7 | 410 | 208 | 50.7 |
| 2 | 14 | 525 | 10 | 1.9 |
| 2 | 21 | 150 | 0 | 0.0 |
| 3 | 0 | 506 | 418 | 82.6 |
| 3 | 7 | 536 | 187 | 34.9 |
| 3 | 14 | 615 | 19 | 3.1 |
| 3 | 21 | 465 | 0 | 0.0 |

**Table S4.** *Anopheles stephensi* and *Aedes aegypti* egg viability post desiccation comparison at 15°C

| **Mosquito strain** | **Replicate** | **Post-desiccation hatching timepoint (day)** | **Egg count** | **Larvae count (L4)** | **Larval survival (%)** |
|---|---|---|---|---|---|
| LVP-IB12 | 1 | 0 | 191 | 153 | 80.1 |
| LVP-IB12 | 2 | 0 | 233 | 202 | 86.7 |
| SDA500 | 1 | 0 | 128 | 24 | 18.8 |
| SDA500 | 2 | 0 | 266 | 34 | 12.8 |
| STE2 | 1 | 0 | 344 | 88 | 25.6 |
| STE2 | 2 | 0 | 202 | 32 | 15.8 |
| LVP-IB12 | 1 | 7 | 303 | 256 | 84.5 |
| LVP-IB12 | 2 | 7 | 177 | 158 | 89.3 |
| SDA500 | 1 | 7 | 298 | 0 | 0 |
| SDA500 | 2 | 7 | 306 | 0 | 0 |
| STE2 | 1 | 7 | 212 | 0 | 0 |
| STE2 | 2 | 7 | 349 | 0 | 0 |
